# Polyimide-Based Flexible Multi-Electrode Arrays for Epilepsy: Synthesis, Microfabrication, and In Vivo Validation In A Rodent Model of Epilepsy

**DOI:** 10.1101/2023.09.12.557325

**Authors:** Kshitij Kumar, Kaustubh Deshpande, Naveen Kalur, Garima Chauhan, Deepti Chugh, Subramaniam Ganesh, Arjun Ramakrishnan

**Affiliations:** Department of Biological Sciences & Bioengineering, Mehta Family Centre for Engineering in Medicine, Indian Institute of Technology Kanpur, Uttar Pradesh, India, 208016; Eywa Neuro Pvt. Ltd. Mumbai, Maharashtra, India, 400086

**Keywords:** Neural Interface, Thin films, Biocompatibility, Epilepsy Monitoring, Electrocorticography (ECoG) arrays, Electrochemical Impedance Spectroscopy

## Abstract

Neurological disorders such as epilepsy, Parkinson’s disease, are rising globally, with conditions like drug-resistant epilepsy affecting millions of patients for whom traditional pharmacological treatments are ineffective. Implantable neural devices have shown great promise in managing these conditions, but their accessibility is limited due to high costs and the availability of suitable biocompatible materials.Thin film implantable neural interfaces hold immense promise over conventional clinical electrodes, offering higher resolution, flexibility, and improved integration with neural tissue. However, their widespread use, especially for flexible interfaces, is limited by the lack of customizable and medical grade materials. We report a novel synthesis method for *ISO 10993-11* compliant polyamic acid that enables the fabrication of biocompatible polyimide films tailored for neural implants. Using this material, we developed 4 and 32 channel depth and surface electrodes, including custom whole brain ECoG arrays. These were implanted in the laforin knockout mice, a validated model of drug-resistant epilepsy, to monitor spontaneous seizures. Both acute and 12 day recordings demonstrated mechanical flexibility, long term stability, and excellent biocompatibility. This study presents a clinically safe material platform and a complete fabrication pathway for building thin film neural interfaces, paving the way for broader clinical use in applications such as epilepsy monitoring and stereo EEG.

## 1. INTRODUCTION

The global incidence of neurological disorders and mental health concerns have been rising rapidly, with conditions like epilepsy, Parkinson’s disease, and depression affecting millions worldwide. The World Health Organization (WHO) reports that neurological disorders account for nearly 6.3% of the global disease burden, with epilepsy alone affecting around 50 million people globally. Similarly, mental health disorders, including depression, have become leading causes of disability, significantly contributing to the global health burden (WHO, 2021). While pharmacological treatments remain the primary therapy, a substantial percentage of patients—up to 30% in the case of epilepsy—are resistant to drug treatments (Kwan & Brodie, 2000). In these drug-resistant cases, implantable neural devices, such as deep brain stimulation systems, vagus nerve stimulators (VNS), and cortical implants, have demonstrated clinical success in providing therapeutic relief.

These devices have been particularly effective in managing conditions such as epilepsy, depression, and Parkinson’s disease, especially when traditional pharmacological therapies fail. For instance, DBS has been widely adopted for treating Parkinson’s disease, while responsive neurostimulation (RNS) has proven to be an effective solution for epilepsy patients with drug-resistant seizures (Fisher et al., 2010). However, the cost of these procedures and devices places them beyond the reach of many, particularly in low- and middle-income countries (LMIC) where healthcare resources are limited. This underlines the urgent need for affordable, scalable, and reliable neural implants that can broaden access to these life-saving technologies.

MEMS-based thin film technology offers promising solutions to some of these challenges. It has driven advancements in neural interfaces by enabling device miniaturization and the development of high-channel-count arrays that are less invasive to brain tissue. Traditional materials like silicon and metal microwires, although providing excellent signal-to-noise ratios, have drawbacks such as stiffness, brittleness, and biocompatibility concerns, limiting their broad clinical use (Nicolelis et al., 1997; Seymour et al., 2017). In contrast, flexible substrates like polyimide, parylene, and liquid crystal polymer align more closely with the mechanical properties of brain tissue, reducing foreign body reactions and improving long-term stability (Weltman et al., 2016; Wang et al., 2020). Polyimide is especially notable for its dielectric properties, flexibility, and thin-film fabrication potential, making it a popular choice for neural interfaces (Constantin et al., 2019). However, the limited availability of medical-grade polyimide precursors continues to hinder the widespread clinical adoption of MEMS-based thin film neural interfaces for acute neurosurgical applications.

Polyimide thin films are synthesized from polyamic acid, which undergoes spin coating and high-temperature curing to convert into polyimide through the imidization process (Chung et al., 2017). Commercially available polyamic acids, such as Pyralin 2610/2611 by HD Microsystems and U-Varnish by UBE Industries, are commonly used in electronic applications but are generally not approved for human medical use (Hassler et al., 2010). The specific properties of polyimide, determined by the monomers used in its synthesis, particularly biphenyl dianhydride (BPDA) and p-phenylenediamine (p-PDA), are crucial for its biocompatibility and stability (Constantin et al., 2019). Despite its potential, the scarcity of medical-grade polyimide limits its use in human clinical settings.

One of the critical applications of neural electrodes is in epilepsy monitoring, where stereotactic electroencephalography (sEEG) and depth electrodes are used to detect and monitor seizures in patients with drug-resistant epilepsy. Accurate neural recordings from depth electrodes are essential for identifying epileptic foci and guiding surgical interventions. Flexible and biocompatible electrodes, such as those made from polyimide, offer significant advantages, as they reduce tissue damage and foreign body responses during long-term implantations. Extended monitoring in animal models, like the laforin knockout mice, provides valuable insights into disease progression and potential treatments (Liu et al., 2019; Kullmann et al., 2022).

In this study, we address the critical limitation in translating MEMS-based microelectrode technologies to acute neurosurgical interventions like microelectrode recordings (MER) and sEEG. By developing a custom polyamic acid chemistry, we demonstrate its application in the microfabrication of neural interfaces, enabling MEMS technology’s full potential, such as channel scaling and flexibility for human neurosurgical applications, particularly in epilepsy monitoring. The following objectives were achieved: 1. Synthesis of BPDA polyimide. 2. Characterization of surface roughness, thermal stability, chemical inertness, and insulation properties of the polyimide film. 3. Microfabrication of 4 to 32-channel, 10 µm thick, depth and surface neural microelectrode arrays. 4. Benchtop electrochemical characterization of the electrode-electrolyte interface. 5. Passing ISO 10993-11 acute systemic toxicity tests for synthesized polyimide thin films. 6. Surface and intracortical implantation of the electrode arrays. 7. Demonstration of high-quality neural activity during acute surgeries, including isolated single units and quantified signal-to-noise ratios in a healthy control as well as a rodent model of epilepsy (laforin knockout). 8. Long term implantation of tetrodes (up to 12 days) in the laforin knockout mice to register epileptic seizures.

## 2. MATERIALS AND METHODS

### 2.1. Polyamic Acid Synthesis

Polyamic acid was synthesized through an equimolar polycondensation reaction between s-BPDA (Merck, India) and p-PPD (Merck, India) in dimethylacetamide (DMAc) (Leo Chemo Plast Pvt. Ltd, India). The reaction was carried out in a 500 ml four-necked round-bottom flask fitted with a water condenser, nitrogen purge, agitator, and thermocouple. Before introducing any reagents, the flask was thoroughly purged with nitrogen gas to ensure an inert environment.

The synthesis began by adding 250 g of DMAc to the flask, which was heated to 30 °C. Then, 0.078 moles of p-PPD were added to the DMAc, and the solution was stirred while being heated to 90 °C for two hours, allowing complete dissolution. After cooling to 50 °C, an equimolar amount of s-BPDA was introduced, and the mixture was stirred continuously for three hours. The temperature was then reduced below 30 °C, and the mixture was stirred for an additional 7–8 hours until a viscous yellow solution formed. To remove unreacted particles, the polyamic acid was filtered using a Whatman Grade 41 filter paper and Buchner funnel under vacuum. The final product was degassed under vacuum for 24–48 hours to eliminate entrapped gases.

### 2.2. Material Characterization

Thermal and curing properties of the synthesized polyamic acid were characterized using thermogravimetric analysis (TGA) and differential scanning calorimetry (DSC), based on three independent samples (N = 3). TGA was used to assess mass changes in response to a controlled temperature increase, providing insights into thermal stability and curing behavior. DSC measured the energy required to raise the temperature of the polyamic acid, indicating the imidization point. These analyses guided the development of a curing process aimed at achieving a high degree of imidization.

The synthesis and curing process was further confirmed using Fourier Transform Infrared Spectroscopy (FTIR) on five polyamic acid and cured polyimide samples (N = 5). The presence of characteristic N-H, C=C, and C=O peaks in the FTIR spectra verified successful synthesis and imidization. Additionally, TGA was used to re-evaluate polyamic acid under verified curing conditions, with minimal weight change indicating complete imidization as the solvent (DMAc) and H₂O were released.

To assess chemical inertness, test structures were fabricated by encapsulating metal lines with polyimide. If the polyimide was compromised, a drop in impedance was observed when tested with electrochemical impedance spectroscopy (EIS) in saline. These test structures were also exposed to various cleanroom chemicals to determine chemical resistance. Surface quality of polyimide films was evaluated using atomic force microscopy (AFM), scanning for surface roughness and pinholes across multiple samples.

Moisture absorption, critical for neural interface longevity, was assessed using TGA. Polyimide films were soaked in deionized water for 24 hours, dried with nitrogen gas, and then heated at 100 °C for 10 minutes. Moisture uptake was calculated as the percentage increase in weight relative to the initial dry weight.

### 2.3. Microfabrication Process

The microelectrodes were fabricated using a two-mask process. Initially, polyamic acid was spin-coated onto RCA-cleaned silicon wafers at 5000 rpm. The wafers were soft-baked at 100 °C for 5 minutes, followed by 120 °C for another 5 minutes. Curing to convert the polyamic acid to polyimide was achieved in a nitrogen-filled furnace at 350 °C. The cured polyimide was roughened using O₂ plasma (PDC-32G-2, Harrick Plasma Systems) to enhance adhesion for subsequent layers.

Next, a bilayer lift-off pattern was created using S1813 photoresist (Microresist Technology) and LOR3B lift-off resist (Kayaku Advanced Materials). A metal stack comprising Titanium (20 nm), Platinum (50 nm), Gold (200 nm), Platinum (50 nm), and Titanium (20 nm) was deposited using a multi-target sputtering system (AJA Systems). Lift-off in Remover PG defined the electrode traces, pads, and bondpads, with a critical dimension of 7µm. Following this, the wafers underwent O₂ plasma descum to remove any residual photoresist.

A second polyimide layer was spin-coated and cured similarly to achieve a total polyimide thickness of 10 µm. Electrode patterns and bondpads were defined using an aluminum (Al) hard mask. 100 nm of Al was deposited by sputtering, patterned using S1813 photoresist, and wet-etched in Type A aluminum etchant (Transense). O₂ plasma was used to etch the polyimide, creating the final device structure by selectively opening the electrodes and bondpads. The Al mask was then removed, and the devices were released from the wafer by soaking in isopropanol and peeling them off using fine-tipped tweezers.

### 2.4. Impedance Characterization

Impedance of the fabricated microelectrodes was characterized using electrochemical impedance spectroscopy (EIS) with a Metrohm Potentiostat (STAT-I-400S). A three-electrode setup was used, comprising a silver/silver chloride reference electrode, a 0.5 mm diameter platinum wire counter electrode, and 1X phosphate-buffered saline (PBS) as the electrolyte to mimic cerebrospinal fluid. Impedance measurements were taken from 100 Hz to 10 kHz, applying a 50 mV sinusoidal signal with three points per decade.

### 2.5. Surgical Protocol and Animal Preparation

Surgical procedures were approved by the Institutional Animal Ethics Committee (IAEC project number: IITK/IAEC/2021/1137). C57BL6 mice and transgenic C57BL6 laforin knockout mice were used across multiple recording sessions. To stiffen the flexible electrode tip before implantation, tetrode depth probes were dip-coated with polyethylene glycol (PEG, P2906, Sigma Aldrich, USA) melting point (50 °C-60 °C.

Mice were anesthetized with isoflurane (5% induction, 1–1.5% maintenance) in 100% oxygen. Anesthesia depth was monitored using the toe pinch withdrawal reflex and re-evaluated every 10–30 minutes. A heating pad was used for thermoregulation. Lidocaine (0.1 mg/kg) was administered subcutaneously for local anesthesia. A primary incision was made along the midline, and the bregma was located using a surgical microscope. A 1 mm × 1 mm craniotomy was performed using a micro-drill, taking care not to damage the dura or blood vessels.

Surface area, dimensions and implant location of the neural probes: A 5 mm long, 4-channel polyimide depth probe (400 µm² (20 x 20 µm) platinum sites) was implanted into the somatosensory cortex (coordinates: AP: -0.9, ML: +3, DV: -1.5) (Franklin and Paxinos (2007)). An ECoG array (7.5 mm long, 4 electrodes, 6400 µm² (800 x 800 µm) electrode area) was placed over the left cortex, guided by a robotic stereotaxic arm (Neurostar, GMBH, Germany) using the mouse brain atlas (Fig. 4). Similarly for the whole brain ECoG (32 Channel custom probe) recording an surface array of dimensions (8.5 mm long, 3.3mm width of each flap, 32 electrodes, 500 µm diameter of the electrode) was placed carefully on the brain covering both hemispheres of the animal’s brain ( Fig. 8b).

For semi-chronic recordings, probes were implanted for durations of up to 12 days, with electrophysiological recordings carried out for up to 10 days post-implantation. Daily recording sessions were carried out following the chronic neural probe implantation procedure, and a 3-day recovery period.

### 2.6. Electrophysiological Recordings

Electrophysiological data were acquired using the Intan RHS stim/recording system (Intan Technologies), with recordings conducted in a custom Faraday cage to prevent signal loss. The Intan RHS32 low-noise amplifier headstage was used, with signals digitized at 30 kHz. Real-time visualization was aided by applying a 50 Hz notch filter to remove electrical noise and a bandpass filter (300–6000 Hz) for neuronal activity monitoring.

### 2.7. Neural Data Analysis

Neural data were analyzed using the **SpikeInterface** library (v0.100) in Python (v3.10). Wideband neuronal signals were re-referenced using central median referencing and bandpass-filtered between 300–6000 Hz. Prior to spike sorting, signals were Z-score normalized and whitened to mitigate artifacts. Spike sorting was performed using the **Mountainsort5** (MS5, Scheme 2) algorithm, which was trained and tested on the entire duration of each recording session.

To assess Single Unit Activity (SUA), the interspike interval (ISI) ratio of the clusters was used. Clusters exhibiting an ISI ratio < 0.5 were classified as SUA, whereas for multiunit activity (MUA) all clusters were used. For ECoG recordings, only MUA was considered for further analysis.

Each recording day may include several approximately 30-60 mins long sessions. Number of neuronal clusters were determined for every session for the day and the maximum from a session was reported as the number of neurons for the day. For Signal to Noise ratio (SNR), the average SNR was determined for every session and then the grand average across sessions for the day was determined.

## 3. RESULTS

### 3.1. Polyimide Synthesis, Characterization, and Microfabrication of Neural Probes

The synthesized polyamic acid had an inherent viscosity of 2.0 dL/g, measured using an Ubbelohde viscometer, indicating the desired polymer chain length and solution properties. For film property evaluation, the polyamic acid was spin-coated onto a silicon wafer, followed by soft baking to form a uniform film. As shown in Fig. 2a., optical microscopy confirmed the absence of pinholes or contaminants, demonstrating that the polyamic acid did not outgas during spin coating or soft baking. This suggests that the polyamic acid formulation is stable and can be further diluted with DMAc to produce thinner polyimide layers, aiding in the miniaturization of neural interfaces.

**Fig 1.**
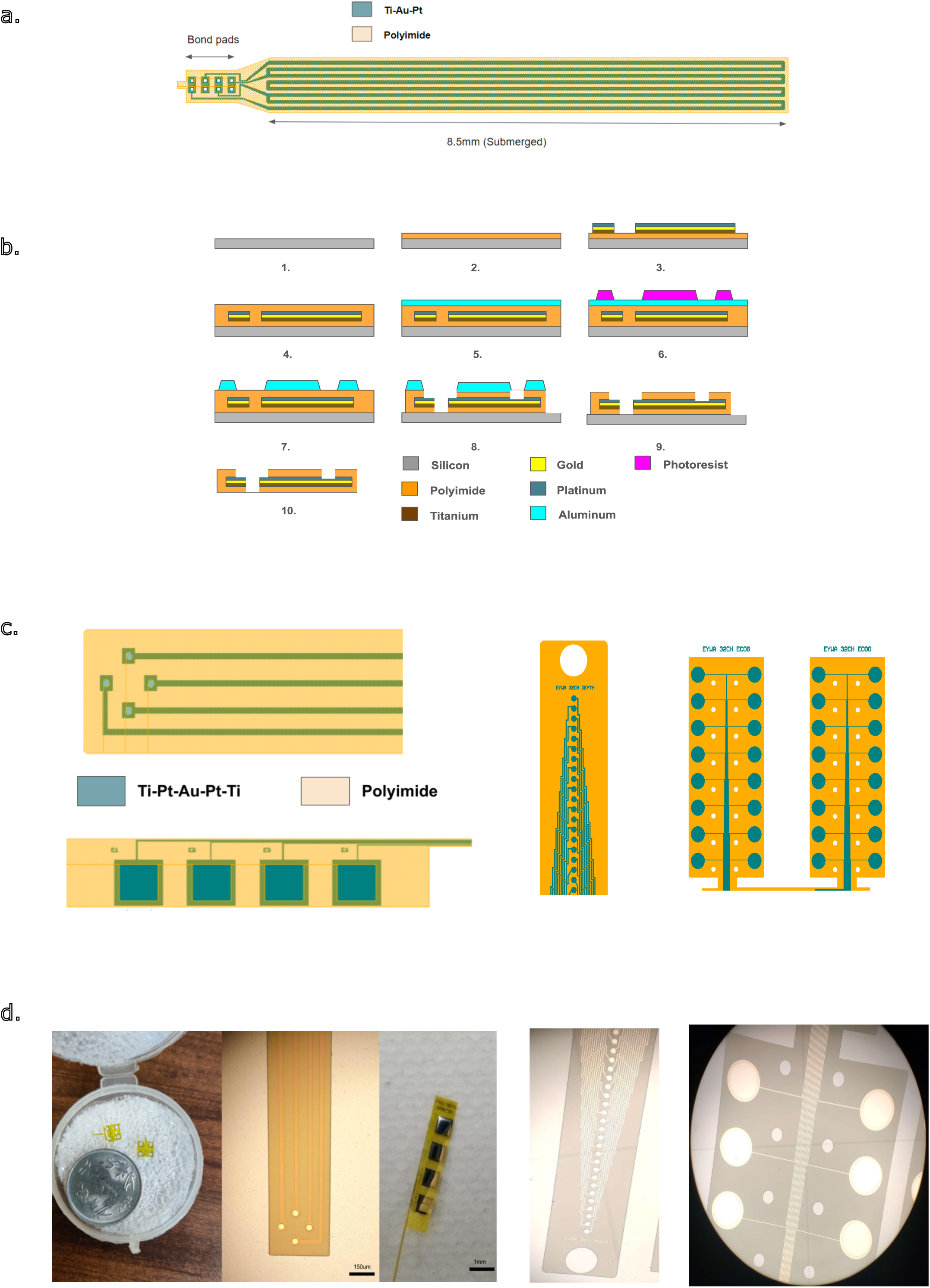
Microfabrication Process and Design Specifications of Polyimide-Based Neural Recording Arrays. a. Shows a test structure for corrosion testing of the polyimide. An 8.5 mm long gold channel arranged in a serpentine manner was encapsulated in 5 µm polyimide. EIS analysis was performed in PBS 1X and the impedance of the test structure was measured to determine whether or not the polyimide thin film was compromised. b. Polyimide MEA Microfabrication Process Steps 1.) RCA clean silicon wafer 2.) Spin coat and cure a base layer of polyimide 3.) Liftoff Ti-Pt-Au-Pt-Ti channels 4.) Spin coat and cure a second layer of polyimide 5.) Evaporate 100 nm Al film 6.) Pattern photoresist over Al layer 7.) Wet etch Al layer to make the hard mask for subsequent polyimide etching 8.) O_2_ plasma etching of polyimide to define outline, electrodes and bond pads 9.)Blanket wet etching of Al hard mask 10.) Release in IPA. c. Schematic representation of the depth and surface ECoG neural arrays, detailing the design, dimensions, and layout of both 32-channel and 4-channel electrode probes. d. Images and close-up views of the actual depth probes (images 1-2 from the left) and surface ECoG arrays (center image), and 32-channel depth array (not used in this study) showcasing electrode design and structural details.

**Fig 2.**
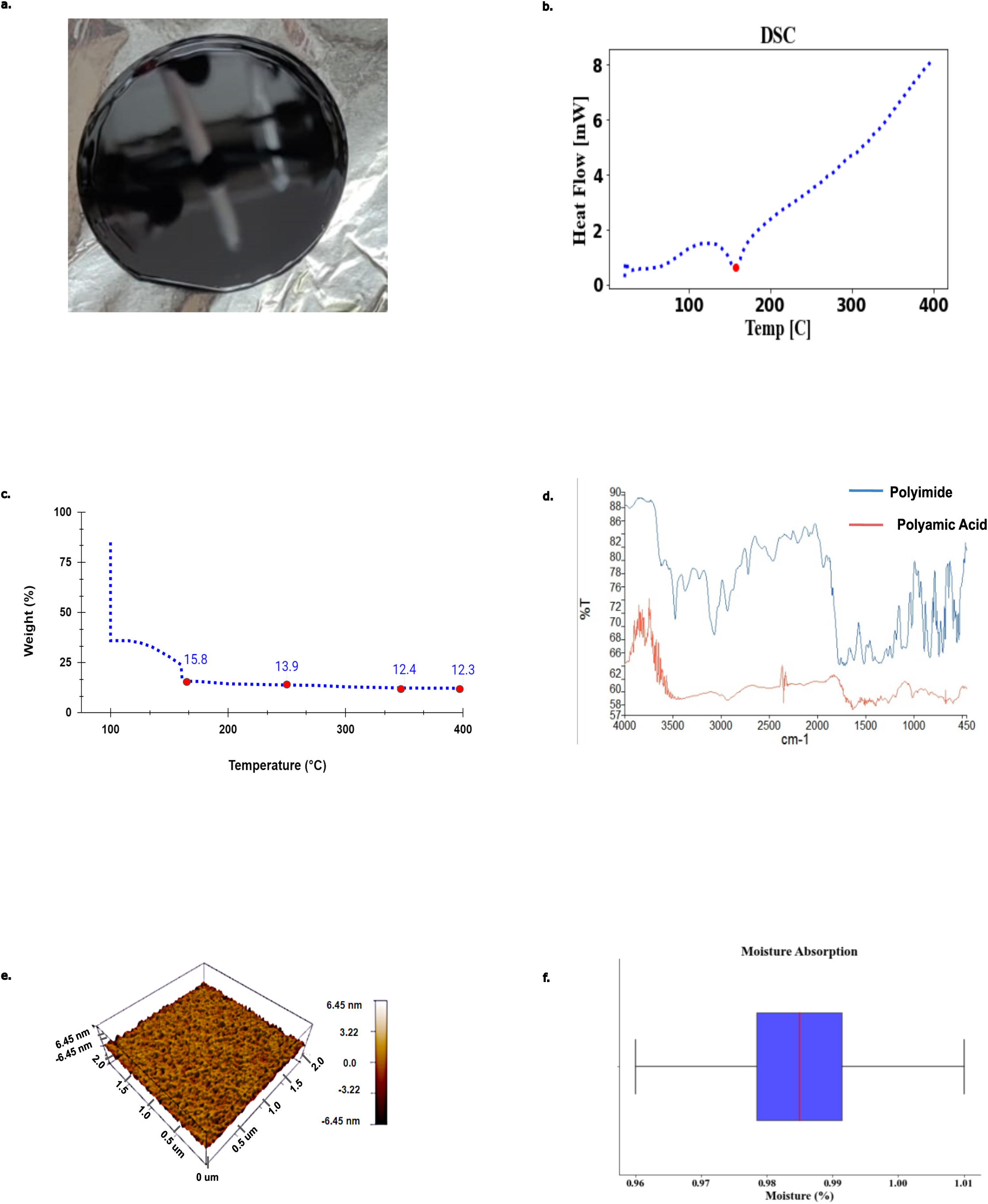
Characterization and Analysis of Polyimide Film Properties. a. Polyamic acid was spin-coated on 50 mm silicon wafers and soft-baked on a hotplate at 100 °C for 5 minutes followed by 120 °C for 5 minutes. The spun wafers were then inspected under high magnification on an optical microscope for the presence of any pinholes or particles. b. DSC Exotherm on a polyamic sample showing non-reversing heat flow behaviour with only a single transition change at 160 °C. The temperature was ramped from 75 °C to 400 °C at 10 °C /min. c. TGA Analysis of the polyamic acid shows the % of weight loss in the polyamic acid sample as it is cured from 75 °C to 400 °C. Most of the weight loss occurs by 160 °C which is the evaporation point of the DMAc solvent. Imdization and subsequent loss of H2O continues up to 400 °C with most of the imidization completed by 350 °C d. Shows the Fourier Transform Infrared (FTIR) spectra for the Polyimide and the polyamic acid. e. Surface roughness distribution of a 2 mm x 2 mm Polyimide film having a film thickness of 5 µm measured using atomic force microscopy. The average surface roughness was around 0.5 nm and the roughness was fairly uniform across the whole cross-section. f. Box plot showing the moisture absorption ratio of N = 6 samples with a mean percentage of 0.984 and standard deviation of 0.016.

Differential scanning calorimetry (DSC) was conducted to examine the thermal behavior of the polyamic acid. The exothermic curve obtained during heating to 400 °C at a rate of 10 °C/min is presented in Fig. 2b. A major thermal transition was observed around 160 °C, corresponding to DMAc evaporation from the polyamic acid. The absence of additional significant transitions and non-reversing heat flow up to 400°C suggest that the processes of cyclization, imidization, and solvent loss continue steadily to this temperature. These results confirm the polyamic acid’s potential for use in high-temperature microfabrication processes.

Thermal stability and imidization behavior of the polyamic acid were further evaluated using TGA (NEXTA STA300, Hitachi). As shown in Fig. 2c., the sample was heated up to 400 °C with a ramp rate of 4 °C/min and isothermal holds at 150 °C, 250 °C, 350 °C, and 400 °C, each for 30 minutes. Continuous weight loss was observed from 160 °C to 400 °C, with residual weights of 13.9% at 250 °C, 12.4% at 350 °C, and 12.3% at 400 °C, indicating the release of water during cyclization and imidization. Beyond 350 °C, the weight change was negligible, confirming that imidization was largely complete. A curing process was adopted with isothermal holds at 150 °C, 250 °C, and 350 °C, followed by a cooling step at 5 °C/min to 75 °C. The solid content of the polyamic acid was measured at 12.3%, showing that the cured polyimide can withstand the high temperatures typically encountered in microfabrication.

The chemical inertness of the cured polyimide was assessed by immersing N = 5 test structures in five chemicals: PRS 2000 (JT Baker), Type A Aluminium etchant (Transene), PBS 1X (Merck, India), Hydrochloric acid (JT Baker), IPA (JT Baker). The distal ends (8.5 mm) of the soldered test structures were soaked for 48 hours at room temperature (27 °C). Post-immersion impedance was tested using electrochemical impedance spectroscopy (EIS) from 100 Hz to 100 kHz in PBS 1X. The polyimide insulation remained intact across all samples, with no change in impedance observed. Optical inspection confirmed secure connections, with no signs of breaks or open circuits, verifying the excellent chemical resistance of the polyimide.

FTIR (Vertex 80, Bruker) analysis was performed to identify the chemical species present in the polyamic acid and cured polyimide films. The FTIR transmission spectra in Fig. 2d. show distinct peaks for N-H stretch bonds between 2900 and 3200 cm⁻¹ in the polyamic acid, with the amide I carbonyl stretch observed at 1653 cm⁻¹. The cured polyimide exhibited characteristic imide peaks, including the imide I C=O stretch at 1768 cm⁻¹, C-N stretch at 1378 cm⁻¹, C-H bend at 1123 cm⁻¹, and C=O bend at 736 cm⁻¹. These results align with prior studies (Diaham et al, 2012), confirming successful synthesis and imidization.

Surface roughness of 5 µm thick polyimide films was measured using atomic force microscopy (AFM, MFP3D Origin, Oxford Instruments). Fig. 2e. illustrates surface scans of 20 mm × 20 mm areas, covering samples from four wafers. With a scanning rate of 1 Hz and 256 points per scan, the measured roughness showed a minimum deviation of -1.31 nm and a maximum deviation of 1.6 nm, yielding an average roughness of 561.43 pm across the film surface. The smoothness of the polyimide films (< 1% surface roughness) ensures compatibility with lithography, metal deposition, and etching processes, making them suitable for neural interface fabrication.

The moisture absorption of the polyimide films was evaluated to determine their suitability for long-term neural interface use. Six independently cured polyimide samples (N=6) were tested for water absorption over 24 hours. Fig. 2f. shows a box plot of the moisture uptake distribution, with a mean absorption of approximately 1%. This low moisture uptake indicates that the polyimide remains stable in aqueous environments, preserving its insulation properties.

Microfabricated depth and surface ECoG arrays are shown in Fig. 1. The devices had a measured thickness of 10.2 µm, verified using a surface profilometer (Dektak XT, Bruker). After release from the silicon wafer in isopropanol, the devices were cleaned with Micro-90 (Cole-Parmer, India), and rinsed with isopropanol and deionized water. Then they were dried and tempered at 120 °C for 6 hours. The arrays were straight, with no kinks or distortions, indicating successful fabrication. The through-hole bondpads were then soldered to an electronic interface board with 1.27 mm male pin connectors.

Electrochemical impedance spectroscopy (EIS) was performed to assess the electrode properties of the arrays. Fig. 3a. & 3c. and Fig. 3b. & 3d. display the average impedance magnitude and phase for N = 5 surface ECoG arrays and N = 5 depth arrays. At 1 kHz, the surface ECoG arrays exhibited an impedance of approximately 10 ± 0.71 kΩ, while the depth arrays measured an impedance of 124 ± 15.23 kΩ. The phase plot showed a capacitive behavior at low frequencies, transitioning to resistive behavior at higher frequencies, consistent with the requirements for both multiunit and single-unit analyses.

**Fig 3.**
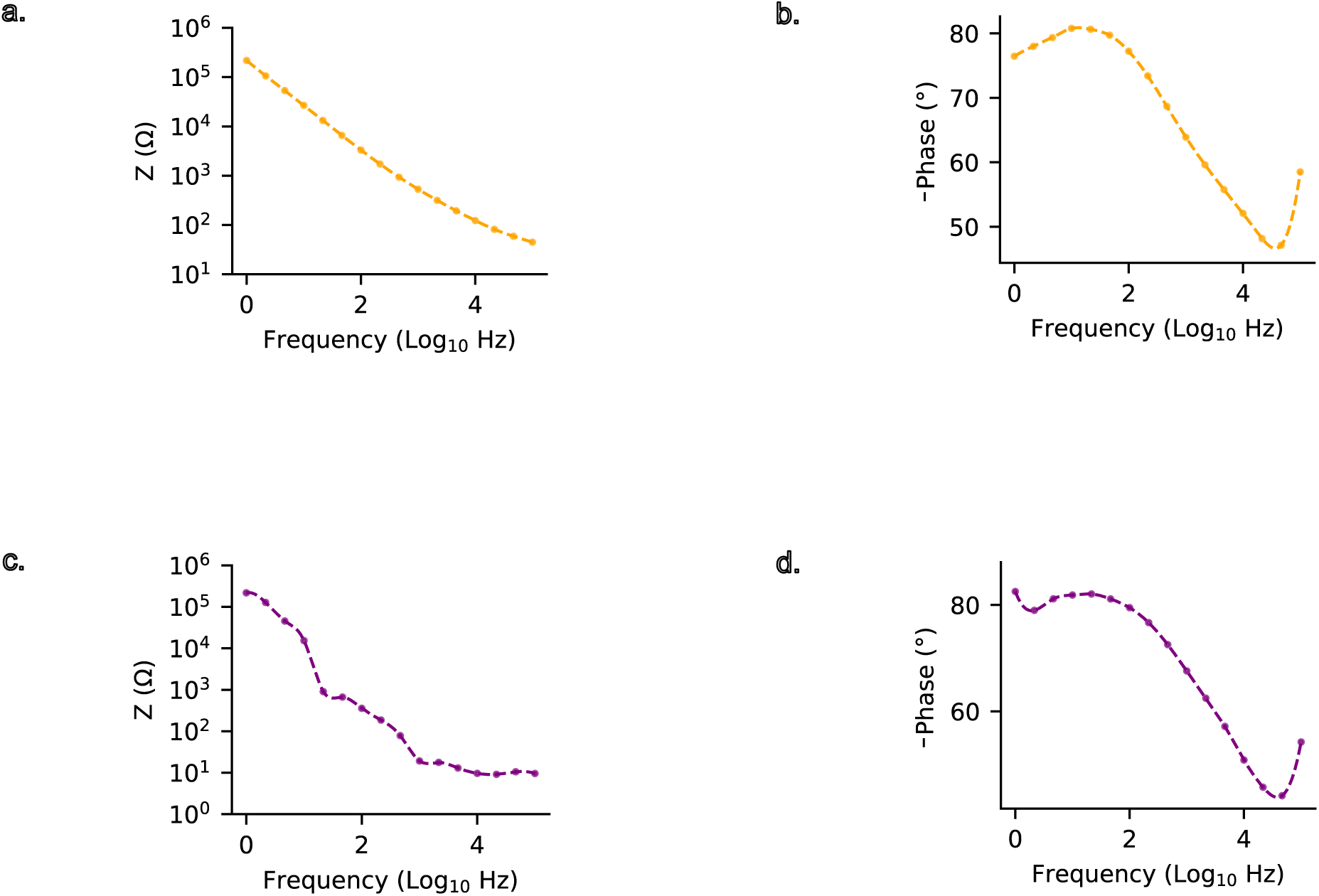
Electrochemical impedance spectroscopy (EIS) measurements of the Depth and Surface Tetrode Arrays. Representative impedance and phase plots are shown between 10 Hz and 10 kHz: Magnitude (a, c) and phase (b,d) of tetrode depth array (a, b) and ECoG (c,d).

Lastly, the polyimide tetrodes were evaluated as per ISO 10993-11 guidelines to assess biocompatibility for medical device applications. Twenty male Swiss Albino mice (Mus musculus; 6–7 weeks; 18.91–20.69 g), housed under CCSEA-approved conditions (21.2–23.1 °C, 53–66% humidity), were randomized into four groups (N = 5/group): Group 1 (polar blank, intravenous), Group 2 (polar test extract, intravenous), Group 3 (non-polar blank, intraperitoneal), and Group 4 (non-polar test extract, intraperitoneal). Dosing at 50 mL/kg followed USP-recommended routes, with extracts confirmed free of particulates post-autoclaving. Clinical observations (mortality, viability, clinical signs) were conducted immediately post-dosing and at 4, 24, 48, and 72 hours, alongside body weight measurements (acclimatization, pre-treatment, pre-necropsy). Necropsy included macroscopic examination of injection sites and organs. No mortality, clinical abnormalities (e.g., lethargy, respiratory distress), or weight loss occurred; all animals exhibited physiological weight gain (e.g., Group 1: +0.5% to +4.1%; Group 4: +0.6% to +2.3%). Gross pathology revealed no abnormalities (NAD in all subjects). The study adhered to OECD GLP principles (QA audits: study plan, item dispensing, extraction, and report phases) and was conducted at an NGCMA-certified facility (RCC Laboratories India), with IAEC approval (Project 309/001). Results confirm the Polyimide’s compliance with ISO 10993-11, demonstrating no acute systemic toxicity at maximal extractable dose volumes, thereby validating its suitability for medical device applications.

### 3.2. Electrophysiological Recordings from Implanted Neural Probes

We implanted different types of electrodes in mice, ranging from 4 channel tetrodes, 4 channel ECoGs (both shown in Fig. 4b.), and 32 channel ECoGs (in Fig. 8 a-b). Data collection lasted either for a single, few hours long, recording session (acute) or for a longer period, up to 10 days long semi-chronic recording session. These recordings were carried out from the somatosensory cortex of the mouse brain. The experiments included both control mice (C57BL/6) and Lafora knockout (LKO), a rodent model that exhibits spontaneous seizures. For the LKO mice, recordings were conducted using multiple configurations: depth 4-channel tetrode recordings (N = 14 for acute experiments), 4-channel ECoG recordings (N=2 for acute experiments), 32-channel ECoG recordings (N=3), and semi-chronic 4-channel depth recordings (N=2). This multi-modal approach allowed for comprehensive characterization of neural activity across different spatial scales and temporal durations.

**Fig 4.**
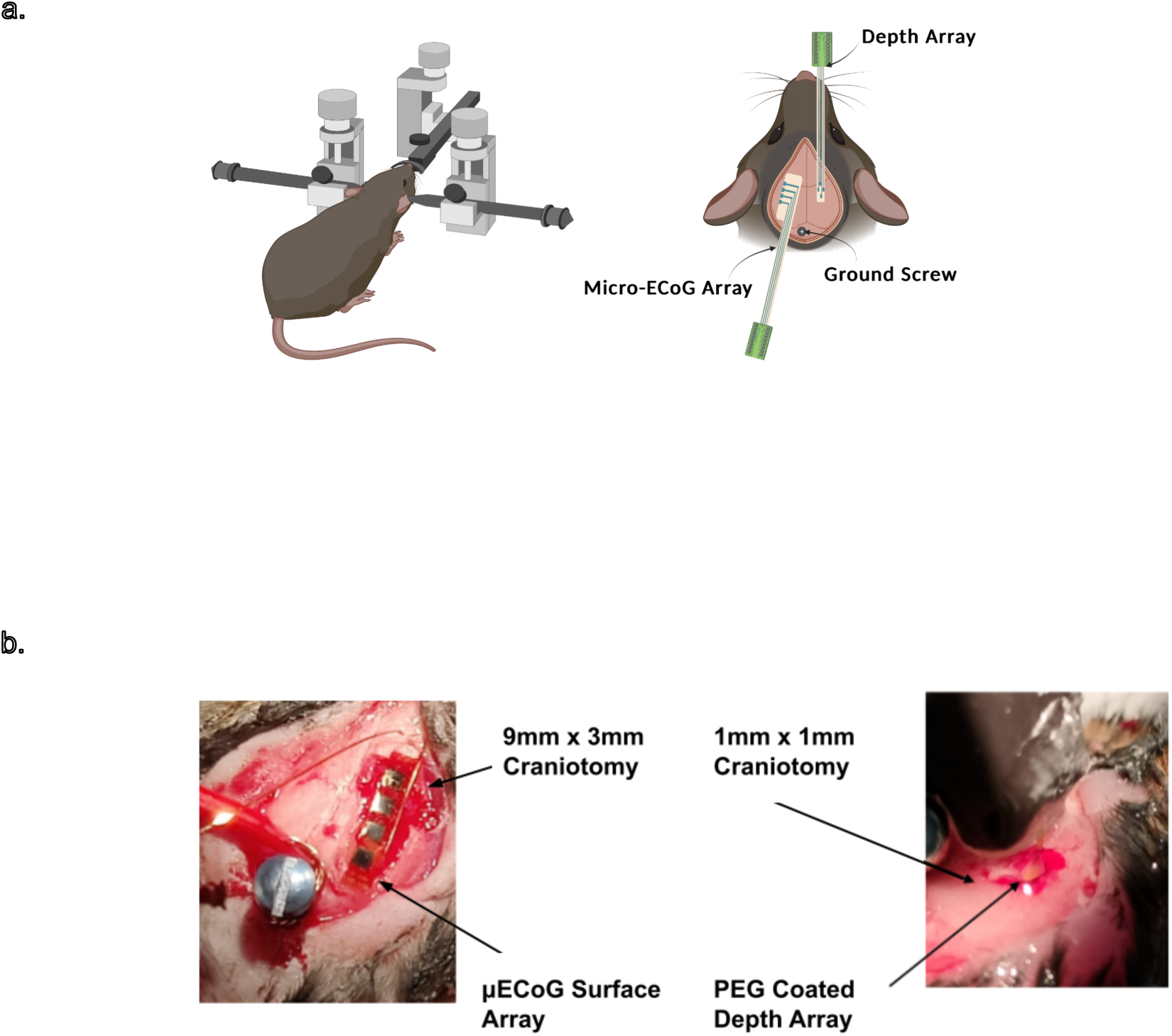
Surgical implantation and positioning of neural recording arrays in a mouse model. a. Illustrative image of an anesthetized mouse mounted on a stereotactic frame, showing the representative placement of the depth tetrode and surface ECoG arrays. b. Intraoperative images demonstrating the surgical placement of microelectrode arrays on the mouse brain, highlighting electrode positioning for neural recordings.

Acute In Vivo neural recordings: In the first set of experiments, we used tetrodes to capture raw neural data, as shown in Figure 5a. Distinct high-frequency spiking activity was observed above the noise seen in the background. The raw data was processed offline to identify single units and local field potentials (MUAs). Action potentials were isolated by first filtering the power in the 300–6000 Hz band. Then using Mountainsort5 the detected action potentials were clustered into putative single units or MUAs (see Methods). Isolated neuronal clusters from the depth probe are depicted in Fig. 5f.

**Fig. 5.**
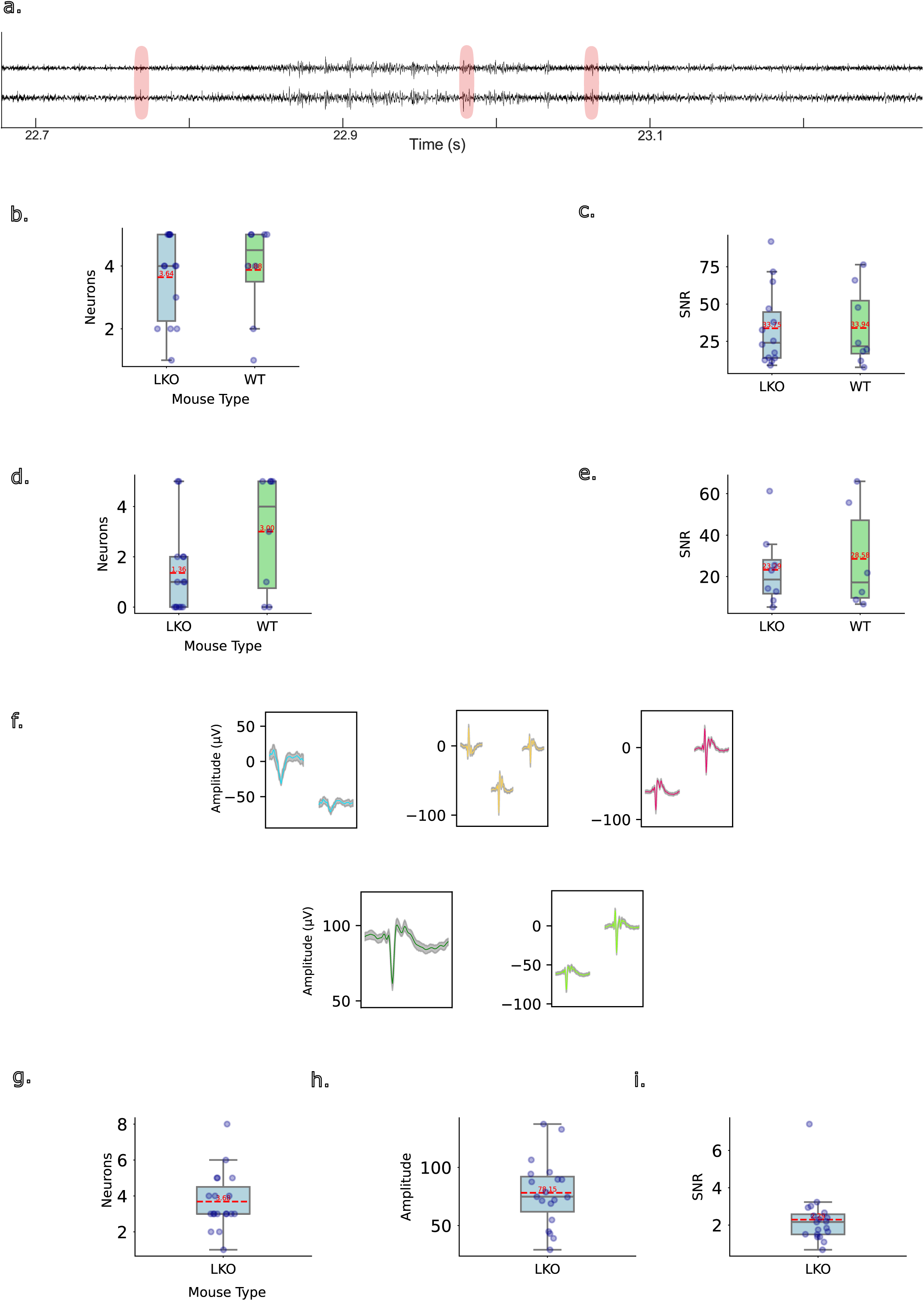
Quality of acute neural recordings from 4-channel depth and surface arrays in wild-type (WT) and LKO mice. Activity based on tetrode depth probe shown in figure panels a-f and 4-ch surface ECoG in panels g-i. a. Representative raw neural traces recorded from the implanted 4-depth probe in an LKO mouse. Two of four channels are shown for reference. Eg: neuronal spikes are highlighted. b. Average multi-unit activity (MUA) recorded from the depth probe across sessions in WT mice (green gradient, N = 8) and LKO mice (blue gradient, N = 14), demonstrating reliable neuronal activity across genotypes. c. Signal-to-noise ratio (SNR) comparison for MUA recordings from the depth probe between WT and LKO mice. d. Average single-unit activity (SUA) from a depth tetrode, capturing consistent cortical population dynamics. e. SNR comparison for SUA recordings from the depth probe between WT and LKO mice. f. Examples of sorted waveforms of SUA recorded during an acute session, showing single-unit isolation across channels. g. Average MUA recorded from the surface (ECoG) probe in LKO mice (N = 2), demonstrating stable population-level activity. h. Average MUA amplitude from the ECoG probe across sessions in LKO mice. i. SNR for surface (ECoG) MUA recordings of LKO mice.

Multi-unit activity (MUA) and single-unit activity (SUA) metrics were assessed across acute recordings obtained using 4-channel tetrode probes. Analysis revealed distinct characteristics between experimental groups LKO and wild type (WT). For MUA recordings (Fig. 5b.), LKO mice yielded 3.64 ± 0.37 neurons per recording with an excellent signal-to-noise ratio (SNR) of 33.75 ± 6.95. These recordings demonstrated optimal presence ratios (1.00 ± 0.00), indicating continuous signal detection throughout the recording period. The mean firing rate was 16.32 ± 3.12 Hz. Similarly, WT mice recordings identified 3.88 ± 0.55 neurons with comparable SNR values (33.94 ± 9.20) and slightly lower but still robust presence ratios (0.96 ± 0.04) and average firing rate of 9.92 ± 3.66 Hz.

Single-unit activity (Fig. 5c.) was successfully isolated from the depth recordings (See Fig. 5f. For template neurons). Group LKO yielded 1.36 ± 0.46 isolated neurons per recording with an SNR of 23.29 ± 6.45 and presence ratio of 0.83 ± 0.12. The mean firing rate for these isolated units was 3.17 ± 1.23 Hz. In contrast, Group WT demonstrated a significantly higher yield of isolated neurons (3.00 ± 0.82) with improved SNR values (28.58 ± 10.49) and presence ratios (0.88 ± 0.08). Firing rates between groups were comparable for SUA (Group W: 3.39 ± 0.82 Hz). These metrics demonstrate the capability of the tetrodes to reliably capture both population-level and single-unit neural dynamics, with distinct electrophysiological signatures between experimental groups.

Signal quality metrics were assessed from multi-unit activity (Fig. 5g.) recorded using 4-channel ECoG in acute preparations (N=2). Analysis focused on characterizing the performance of these surface probes.

Even in the acute ECoG recordings in Group LKO, we observed 4.50 ± 0.50 neurons (MUA) per recording. These units demonstrated a SNR of 4.81 ± 2.61, reflecting the challenges of capturing neural signals from the cortical surface. Despite moderate SNR values, the recordings exhibited optimal presence ratios (1.00 ± 0.00), indicating continuous and reliable signal detection throughout the recording duration. The mean firing rate was 18.31 ± 4.08 Hz, demonstrating consistent neural activity. These metrics support the capability of the 4-channel ECoG tetrode probes to capture neural population activity, though with expected signal quality trade-offs compared to depth recordings.

Semi-chronic In Vivo neural recordings: Neural recordings were evaluated across multiple post-implantation days (days 0, 4, 12 in Fig. 6a.) in two mice to assess long-term signal stability using 4-channel depth tetrode probes. For Mouse 1, signal-to-noise ratio (SNR) values remained stable, with mean 7.89 ± 0.48 on day 0, 8.70 ± 1.49 on day 4, and 8.14 ± 0.88 on day 10. Presence ratios were consistently high, exceeding 0.96 on most days (day 0: 0.98 ± 0.009; day 4: 0.98 ± 0.008; day 10: 0.99 ± 0.011), indicating persistent unit detectability. For Mouse 2, SNR values were also stable over time, with means of 11.00 ± 1.10 on day 0, 17.19 ± 2.87 on day 4, and 7.19 ± 0.05 on day 12, reflecting sustained multiunit signal quality. Presence ratios improved across sessions, from 0.60 ± 0.08 on day 0 and 0.60 ± 0.06 on day 4 to 0.91 ± 0.04 on day 12. Together, these findings highlight the long-term reliability of the probe design in capturing high-fidelity multiunit activity over extended periods in semi-chronically implanted animals.

**Fig 6.**
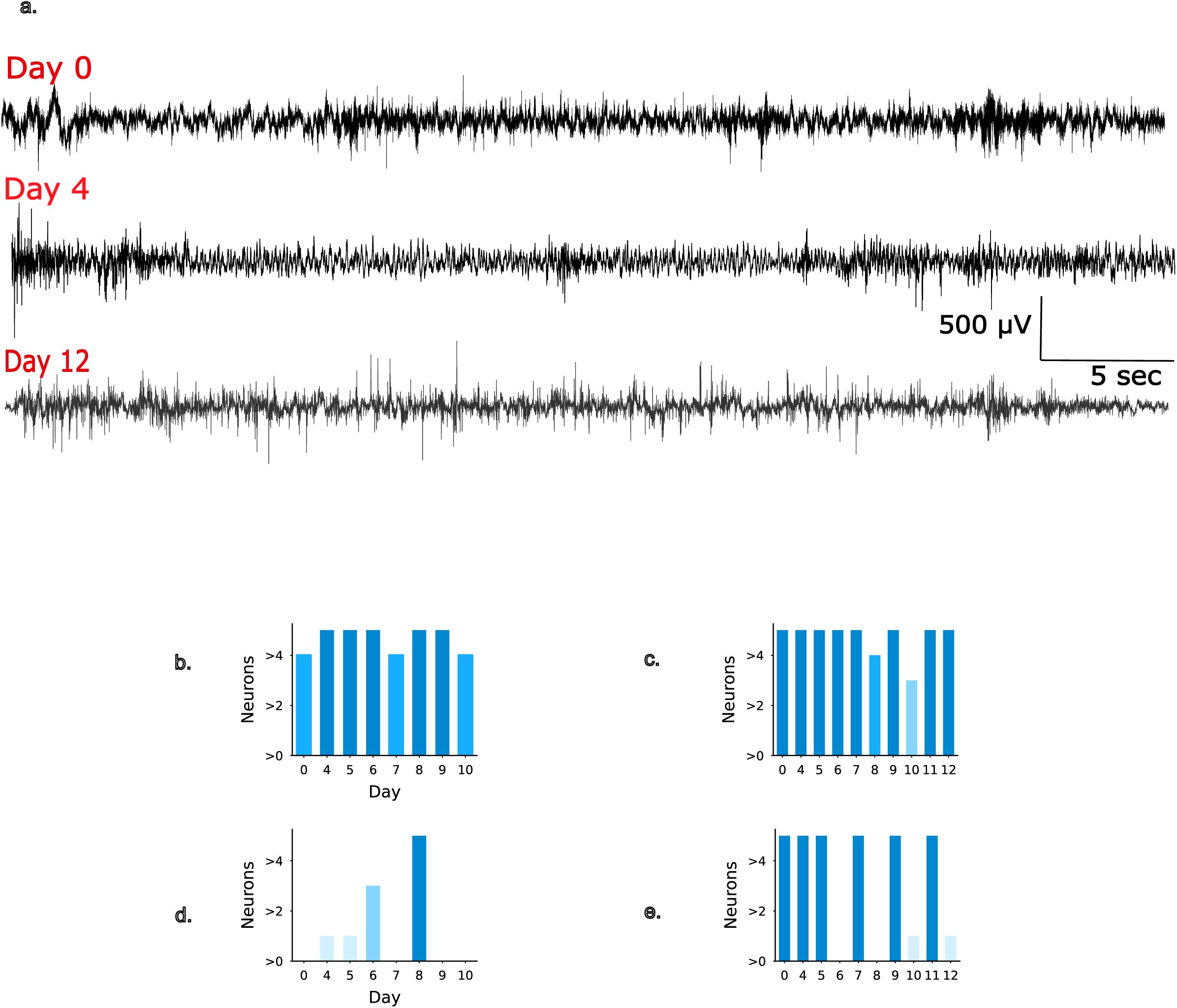
Quality of Longitudinal Neural Recording (up to 12 days) in LKO C57BL/6 Mice. Representative neural signal traces (in a) acquired from semi-chronically implanted depth probes in an LKO C57BL/6 mouse at three distinct time points post-implantation (day 0, day 4, and day 12), demonstrating the change in signal characteristics over the recording period. Quantitative analysis of multiunit activity (b - animal 1, c - animal 2) and single-unit activity (d - animal 1, e - animal 2, following spike-sorting across up to a 12-day implantation period in LKO mice (N=2). Y-axis indicates neuron count, while x-axis represents recording days (day 0 through day 12). Color intensity corresponds to neuronal density, with error bars indicating standard error of mean calculated across multiple ∼1-hour long recording sessions conducted on the respective day.

Out of these MUA, those that passed the SUA criteria were evaluated for their quality, both Mouse 1 (Fig. 6d.) and Mouse 2 (Fig. 6e.) exhibited distinct yet converging trends in signal quality over time in the SUA. Pre-surgery (Day 0), Mouse 2 showed a moderate signal-to-noise ratio (SNR) of 13.24 ± 1.89 with a presence ratio of 0.34 ± 0.06, suggesting decent baseline signal quality, while Mouse 1 had no detectable units at this stage. Following surgery, by Day 4, both animals demonstrated a marked improvement in recording quality—Mouse 2 showed an SNR increase to 20.52 ± 3.58 and stable presence (0.47 ± 0.04), whereas Mouse 1 exhibited a sharp rise in SNR to 55.00 and a presence ratio of 1.00 ± 0.01, indicating strong and reliable unit detection. By Day 8-12, signal quality reduced again: Mouse 1 experienced a drop in SNR to 9.39 ± 0.59 and presence ratio to 0.75 ± 0.06, while Mouse 2 declined to SNR of 7.30 ± 0.00 and a presence ratio of 0.24 ± 0.00 respectively. These patterns likely reflect a combination of mechanical drift, glial scarring, and tissue response over time, which commonly impact semi-chronic electrode recordings.

### 3.3. Epilepsy Monitoring Using Polyimide Probes

To evaluate the effectiveness of our custom-fabricated polyimide-based electrodes for semi-chronic epilepsy monitoring, we implanted tetrode-style probes in the *laforin* knockout (LKO) mice (N = 2) and continuously recorded neural activity over a 7-10 day period (10 day implantation includes 3 days recovery period for mouse 1 and 12 days implantation including 3 days recovery for mouse 2). LKO mice serve as a well-characterized model for spontaneous, recurrent seizures, making them suitable for long-term tracking of epileptiform dynamics.

Recordings from Day 0—immediately post-surgery as anesthesia effects were wearing off—already revealed clear signatures of epileptiform discharges (Fig. 7, Day 0). Representative snippets from Day 4 and Day 12 illustrate both shorter (<10 seconds) ictal discharges and longer (>10 seconds) seizure episodes, which occurred frequently (Fig. 7, Day 4 and 12). These events were observed in both animals and showed strong reproducibility across sessions, underscoring the stability and reliability of the implanted probes in capturing diverse forms of epileptic activity throughout the monitoring window.

**Fig 7.**
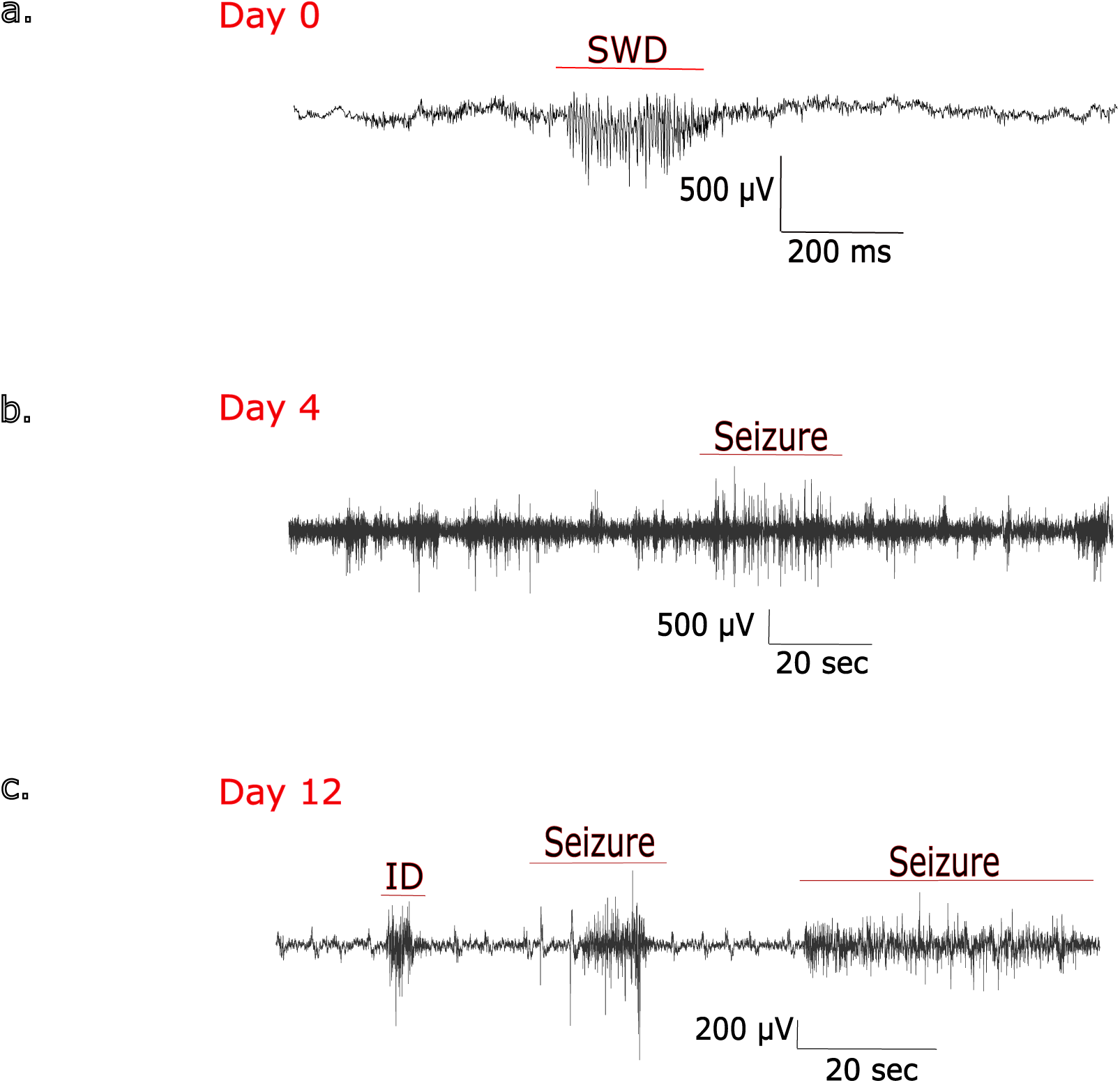
Epileptiform Activity Patterns in LKO C57BL/6 Mice During Semi-chronic Recording. Epileptic features recorded from the depth probe in an LKO mouse of the C57BL/6 strain on day 0 (a), day 4 (b), and day 12 (in c), demonstrating successful implantation and stable recording post surgery to capture epileptiform activity. SWD-spike wave discharge, ID - interictal discharge.

To expand the spatial coverage and improve cortical resolution, we also developed a custom-designed, 32-channel electrocorticography (ECoG) array tailored for high-density surface recordings from the mouse cortex (Fig. 8a-b). This flexible, polyimide-based array features a conformal design that allows it to closely adhere to the cortical surface, ensuring stable contact with minimal tissue disruption and long-term signal fidelity.

**Fig 8.**
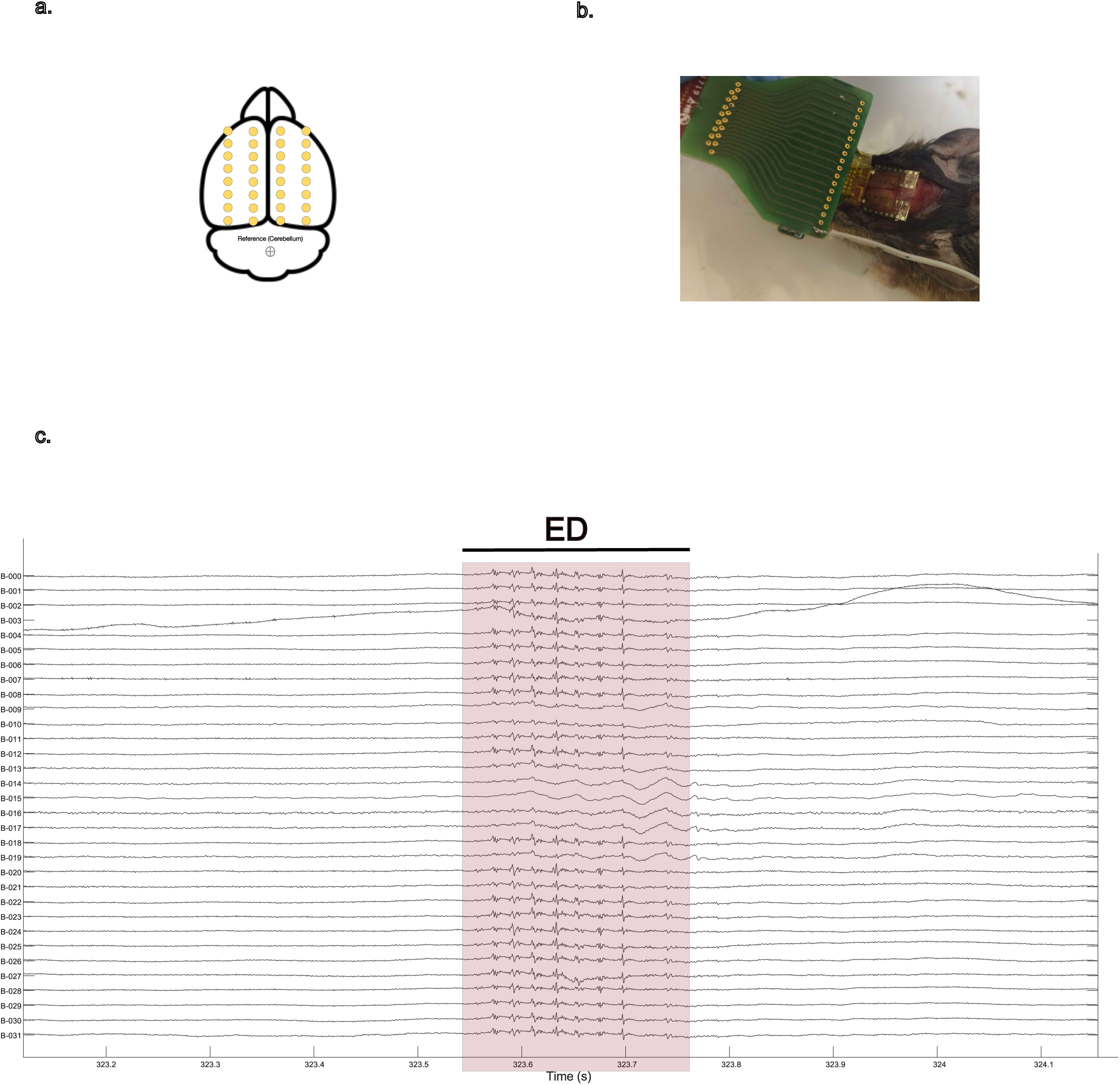
High-Density Surface Mapping of Epileptiform Activity Using Custom 32-Channel ECoG Arrays. a. Layout of the customized 32-channel ECoG electrode array placed on the brain, illustrating the mapping of brain regions and electrode positions designed to identify and isolate the region of interest for epileptiform discharges and the seizure epicenter. b. Shows the superpositioning of ECoG on the mice brain. c. Representative neural traces recorded from the custom 32-channel ECoG array, capturing the transient epileptiform discharge (ED, red box) across the whole brain in C57BL/6 LKO mouse.

The array’s broad spatial footprint supports bilateral recordings across both hemispheres (Fig. 8a.), enabling us to detect and map transient ictal discharges that were present across both cortical hemispheres (Fig. 8c.). These cross-hemispheric seizure events could provide critical insights into the distributed network mechanisms that underlie generalized seizures and interregional synchrony in epilepsy.

Collectively, these results demonstrate that our polyimide-based probes—both depth electrodes and surface ECoG arrays—offer robust performance for semi-chronic epilepsy monitoring. Their ability to record stable, high-resolution neural signals over extended periods, with both temporal and spatial precision, positions them as a valuable tool for probing the functional architecture of epileptic networks and for advancing translational neuroscience research in seizure disorders.

## 4. DISCUSSION

The global burden of neurological and mental health disorders is rising, with epilepsy alone affecting around 50 million people and contributing 6.3% to the global disease burden (WHO, 2021). Up to 30% of patients, particularly those with epilepsy, do not respond to drug treatments (Kwan & Brodie, 2000). For these drug-resistant cases, implantable neural devices like deep brain stimulation (DBS), vagus nerve stimulators (VNS), and cortical implants have shown clinical success, particularly in epilepsy, Parkinson’s disease, and treatment-resistant depression (Fisher et al., 2010). However, their high costs and limited availability in low-income regions emphasize the need for more affordable, scalable, and accessible neural implants.

### 4.1. Need for Custom Polyimide Synthesis

While we acknowledge the decades-long effort in developing polyimide-based electrode arrays, our work addresses a critical and often overlooked gap in the field. Neural interfaces are highly context-specific—device performance depends not only on architecture and implantation site, but also on the mechanical, dielectric, and chemical properties of the materials used, as well as on surgical compatibility. Most existing efforts have relied on proprietary polyimide formulations (e.g., from HD Microsystems or UBE), which limit transparency and hinder customization for neural applications.

In contrast, our study presents an open, end-to-end workflow for the synthesis, characterization, and fabrication of medical-grade polyimide films specifically designed for neural interface use. A key advantage of our in-house approach is the ability to precisely tailor material properties by adjusting the chemistry and processing of the precursor polyamic acid. Parameters such as monomer ratios, solid content, solvent system, and inherent viscosity can be systematically varied to control film thickness, conformality, mechanical flexibility, and thermal stability—properties essential for minimizing tissue damage and enhancing implant longevity.

Moreover, our synthesis process enables modulation of dielectric properties and allows for the incorporation of functional groups or fillers to improve adhesion to metal layers, enhance bio-inertness, or introduce bioactive coatings. This amount of material-level customization is not achievable with commercial polyimides optimized for the electronics industry.

We demonstrate not only the synthesis and material optimization but also the full fabrication and testing pipeline—from spin coating and patterning to in vivo validation of functional neural interfaces in both tetrode and linear configurations. These devices were successfully tested in acute and semi-chronic settings, and our material characterization data support QA/QC standards relevant for translational development. While the electrode designs themselves are conventional, the novelty and impact of our work lie in making a customizable, biocompatible polyimide formulation accessible to the neural engineering community—providing a versatile platform for adapting neural interfaces to specific clinical, anatomical, and surgical needs.

### 4.2. Comparison with Existing Technologies

The impedance values of our fabricated polyimide depth arrays and ECoG probes align with those described by Seymour et al. (2017) and Wellman et al. (2017), suggesting similar electrical properties. Specifically, our tetrode depth arrays (4 channel, electrode area (20 x 20 µm), with inter electrode distance of 150 µm, exhibited an average impedance of 124 ± 15.23 kΩ at 1 kHz, while ECoG arrays (4 channel, electrode area ( 800 x 800 µm), inter electrode distance (1500 µm)) demonstrated an average impedance of 10 ± 0.71 kΩ at 1kHz. These values match the established performance benchmarks for multiunit and single-unit analysis (Han et al., 2024), affirming the effectiveness of our fabrication process while addressing cost limitations.

In terms of neural signal quality, our probes provided reliable single-unit isolation and stable multi-unit recordings, with signal-to-noise ratios (SNR) comparable to other polyimide-based probes, such as those developed by Weltman et al. (2016). Our depth tetrode arrays achieved a SNR 33.75 ± 6.95 in LKO mice and 33.94 ± 9.20 in WT mice, which was higher than that of the surface ECoG probes (4.81 ± 2.61), as expected. Notably, our semi-chronic implants maintained satisfactory signal quality over extended periods, with MUA SNR values remaining above 7.0 even after 10-12 days post-implantation, though some expected signal degradation was observed for SUA recordings. The perfect presence ratios observed in both acute ECoG recordings and early semi-chronic implants further validated the mechanical stability of our design. These results align with prior studies by Horváth et al. (2024), confirming the utility of flexible polyimide thin films in capturing neural activity across a range of frequencies. The synthesis technique and fabrication process for clinical-grade polyamic acid have enabled the production of robust polyimide neural interfaces, positioning them as a cost-efficient alternative to traditional neural probes.

The polyimide-based neural probes developed in this study have demonstrated considerable potential for effective epilepsy monitoring. During in vivo testing, the probes reliably captured critical epileptiform signatures, such as high-frequency oscillations, spiky waveforms, and other seizure-associated discharges. These results align with findings from similar polyimide-based neural interfaces, as reported by Seymour et al. (2017) and Weltman et al. (2016), which also demonstrated effective detection of epileptiform patterns in rodent models. The successful identification of these signals supports the suitability of our probes for accurately monitoring seizure dynamics and localizing epileptic foci, which are crucial for guiding surgical interventions in epilepsy (Staba et al., 2014; Kullmann et al., 2022).

The probes were validated under both acute conditions and over a 12-day implantation period, establishing their durability and consistent performance in extended neural interfacing. This aligns with findings from Liu et al. (2019), who underscored the importance of stable and continuous neural signal acquisition in long-term epilepsy monitoring using flexible thin-film electrodes. Long-term implants are essential for continuous seizure tracking, and our results confirm that the probes can maintain stable, high-quality recordings over prolonged periods. This is also consistent with observations by Rubehn and Stieglitz (2010), who noted the sustained performance of polyimide materials in chronic neural interfaces.

Biocompatibility is critical for neural implants intended for chronic use. In line with this, our polyimide material underwent comprehensive biocompatibility assessments following ISO 10993-11 guidelines at an FDA-certified laboratory (RCC Labs, Hyderabad). These tests confirmed the material’s non-toxicity, minimal inflammation risk, and reduced foreign body response—factors essential for long-term implantation (Ha et al., 2008). The successful results address uncertainties related to material sourcing and ensure compliance with stringent medical device safety standards, enhancing the feasibility of chronic use in epilepsy patients (Wellman et al., 2017). This validation further reinforces polyimide’s role as a reliable material for developing advanced neural interfaces, supporting its application in both acute and long-term epilepsy monitoring.

### 4.3. Broader Implications and Future Work

Our polyimide thin films could potentially find applications beyond acute neurosurgical interventions. With high thermal stability, chemical inertness, and low moisture absorption, these films are suitable for a wide range of biomedical uses, including short-term implants, drug delivery systems, and biocompatible coatings (Ordonez et al., 2012; Diaz-Botia et al., 2022). The material’s versatility supports cross-disciplinary innovation, paving the way for advancements in medical technologies through collaborative efforts. Future research will focus on evaluating the polyimide’s performance in long-term and chronic applications to establish its suitability for continuous neural monitoring and chronic implants (Nicolelis et al., 1997). Additionally, efforts will be directed towards increasing the channel count of fabricated devices to enhance spatial resolution and enable comprehensive mapping of multiple brain regions simultaneously (Buccino et al., 2020). This ongoing research aims to develop advanced neural monitoring solutions, expanding the scope and applicability of this innovative technology in treating neurological disorders.

## ACKNOWLEDGEMENTS

We extend our gratitude to K. Nageswari, Deepti Rukade, and the team at the IIT Bombay Nanofabrication Facility for their support in coordinating schedules for microfabrication. We are also grateful to Jamille Hetke and Rio Vetter at NeuroNexus Technologies for their guidance on polyamic acid and polyimide electrode technologies. Special thanks to the staff at the Central Experimental Animal Facility, IIT Kanpur, for their assistance and expertise in animal surgeries. Deepti Chugh and Garima Chauhan were supported by the Institute Postdoctoral Fellowship, IIT Kanpur, while Kshitij Kumar received support from the ICMR-DHR-CoE. This research was funded by the Indian Council of Medical Research (ITR grant), DBT Wellcome Trust IA intermediate fellowship and IIT Kanpur startup funds awarded to Arjun Ramakrishnan.

## AUTHOR CONTRIBUTIONS

Kaustubh Deshpande (KD) and Arjun Ramakrishnan (AR) conceived the study. KD and Naveen Kalur (NK) were responsible for the design and synthesis of the polyimide and neural probes. Kshitij Kumar (KK), Garima Chauhan (GC), and Deepti Chugh (DC) conducted the animal surgeries and data acquisition. KK analyzed the neural recordings. KK and KD drafted the original manuscript. Subramaniam Ganesh (SG) developed the transgenic mouse model used in this study. All authors reviewed and revised the final manuscript.

